# Phage-antibiotic synergy: cell filamentation is a key driver of successful phage predation

**DOI:** 10.1101/2022.11.28.518157

**Authors:** Julián Bulssico, Irina Papukashvili, Leon Espinosa, Sylvain Gandon, Mireille Ansaldi

## Abstract

Phages are promising tools to fight antibiotic-resistant bacteria, and as for now, phage therapy is essentially performed in combination with antibiotics. Interestingly, combined treatments including phages and a wide range of antibiotics lead to an increased bacterial killing, a phenomenon called phage-antibiotic synergy (PAS), suggesting that antibiotic-induced changes in bacterial physiology alter the dynamics of phage propagation. Using single-phage and single-cell techniques, each step of the lytic cycle of phage HK620 was studied in *E. coli* cultures treated with either ciprofloxacin or cephalexin, two filamentation-inducing antibiotics. In the presence of sublethal doses of antibiotics, multiple stress tolerance and DNA repair pathways are triggered following activation of the SOS response. One of the most notable effects is the inhibition of bacterial division. As a result, a significant fraction of cells forms filaments that stop dividing but have higher rates of mutagenesis. Antibiotic-induced filaments become easy targets for phages due to their enlarged surface areas, as demonstrated by fluorescence microscopy and flow cytometry techniques. Adsorption, infection and lysis occur more often in filamentous cells compared to regular-sized bacteria. In addition, the reduction in bacterial numbers caused by impaired cell division may account for the faster elimination of bacteria during PAS. We developed a mathematical model to capture the interaction between sublethal doses of antibiotics and exposition to phages. This model shows that the induction of filamentation by sublethal doses of antibiotics can amplify the replication of phages and therefore yield PAS. We also use this model to study the consequences of PAS on the emergence of antibiotic resistance. A significant percentage of hyper-mutagenic filamentous bacteria are effectively killed by phages due to their increased susceptibility to infection. As a result, the addition of even a very low number of bacteriophages produced a strong reduction of the mutagenesis rate of the entire bacterial population. We confirm this prediction experimentally using reporters for bacterial DNA repair. Our work highlights the multiple benefits associated with the combination of sublethal doses of antibiotics with bacteriophages.

## Introduction

Antimicrobial resistance (AMR) is one of the major threats in public health, and a recent study estimates that almost 5 million deaths were associated worldwide with AMR in 2019 [1]. Moreover, in the context of the current COVID-19 pandemic, tackling antibiotics resistance remains a high priority for the World Health Organisation (WHO), and the medical community is highly concerned by the increased risk of antibiotics misuse as well as by the decreased concern of worldwide governments to face this major public health issue, which is one of the consequences of the viral pandemic [2,3]. In this context, phage therapy is more than ever regaining scientific and medical interests after several decades of exclusive use of antibiotics. In Europe, since January 2018, phage magistral preparations have been approved and applied in Belgium [4], whereas in others countries, compassionate usage has been approved in combination with different antibiotics [5,6]. Note that a genetically modified bacteriophage has been successfully used recently for the treatment of an immunocompromised cystic fibrosis young patient, dying from a *Mycobacterium abscessus* infection [7]. Interestingly, most recent therapeutic experiences combine the use of antibiotics together with specific phage preparations and proved to be more effective in preclinical studies [8,9].

It’s been recognized that phage bactericidal efficacy increased massively when phages and antibiotics were used in combination, a phenomenon called PAS, for Phage-Antibiotic Synergy [10]. Several laboratories worldwide are studying PAS but most studies are restricted to a limited number of combinations, and rarely focused on the mechanistic elucidation of this effect [11–13]. Interestingly, ecological and epidemiological considerations are emerging nowadays as keys to elucidate and predict the consequences of such combinatorial treatments [14]. Moreover, combinatorial treatments not only increase the killing potential of phages and antibiotics, but can also re-sensitize antibiotic tolerant bacterial strains [15–18]. In addition, this combinatorial approach may also foster the decrease of the antibiotic dosage and reduce the selective pressure on bacteria, thus the potential for antibiotic resistance emergence during therapy [16].

In this work we investigated the link between increased phage infection and antibiotic driven bacterial filamentation caused by ciprofloxacin and cephalexin, two antibiotics that belong to different classes fluoroquinolone and ß-lactam. They both promote cell elongation but through different mechanisms. Our study emphasizes on the need to investigate PAS at the single cell level to elucidate phage susceptibility in the different bacterial subpopulations elicited by the antibiotic treatment. Accordingly, we developed single-cell as well as single-phage approaches to monitor the critical steps of phage infection using the temperate phage HK620 and its host *Escherichia coli* TD2158 PL4 as an experimental model. We show that both antibiotics are promoting increased phage killing in liquid as well as in solid cultures. We developed a mathematical model to study the influence of sublethal doses of antibiotics on phage-bacteria dynamics. This theoretical framework yields predictions on the synergistic effects of antibiotics and phages on bacterial population growth and on the rise of antibiotic resistance. This joint theoretical-experimental approach elucidates the underlying processes leading to PAS and shows that the combination of phages and antibiotics could have an additional value in terms of lowering the influx of antibiotic resistance mutations.

## Results

### Both cephalexin and ciprofloxacin act synergistically with phage HK620

This study addresses the phenomenon of Phage-Antibiotic Synergy, understood as a boost in phage propagation in the presence of antibiotics at low dosages, that otherwise will not lead to significant cell death [10]. PAS measurements were performed in semi-solid and liquid cultures using HK620 dedicated bacterial host *E. coli* TD2158 PL4, which was cured for all resident prophages that proved to be inducible [19,20]. Although HK620 is a temperate phage, it shows a very high spontaneous induction level at 30ºC, and behaves fully lytic at 37ºC [20]. PAS on a semi-solid culture was tested by measuring lysis plaque radii generated by HK620 on a top agar assay (Fig. 1a, b). In the presence of either ciprofloxacin or cephalexin used at ½ of the MIC, phage HK620 average plaque radius increased by 32.8% (p < 0.001) and 9.6% (p<0.01), respectively. PAS was also measured on liquid cultures by comparing bacterial OD_600_ reduction upon phage infection in the presence or absence of the same antibiotic concentrations as above (Fig. 1c). After two hours of incubation with or without antibiotic, the same number of phages HK620 phages was added. In the presence of any of the mentioned antibiotics, bacterial killing occurred significantly faster and more efficiently than in the untreated culture. The presence of the antibiotics at sub-lethal concentrations did not impair culture growth, as observed by OD_600_ measurements in the absence of phage. These results confirm the synergistic behaviour of phage HK620 in combination with ciprofloxacin or cephalexin used at half of the Minimal Inhibitory Concentration (MIC) concentrations: the presence of sublethal antibiotic concentrations enhanced phage propagation and killing of *E. coli* TD2158 PL4 in liquid and semi-solid media.

**Figure 1:**
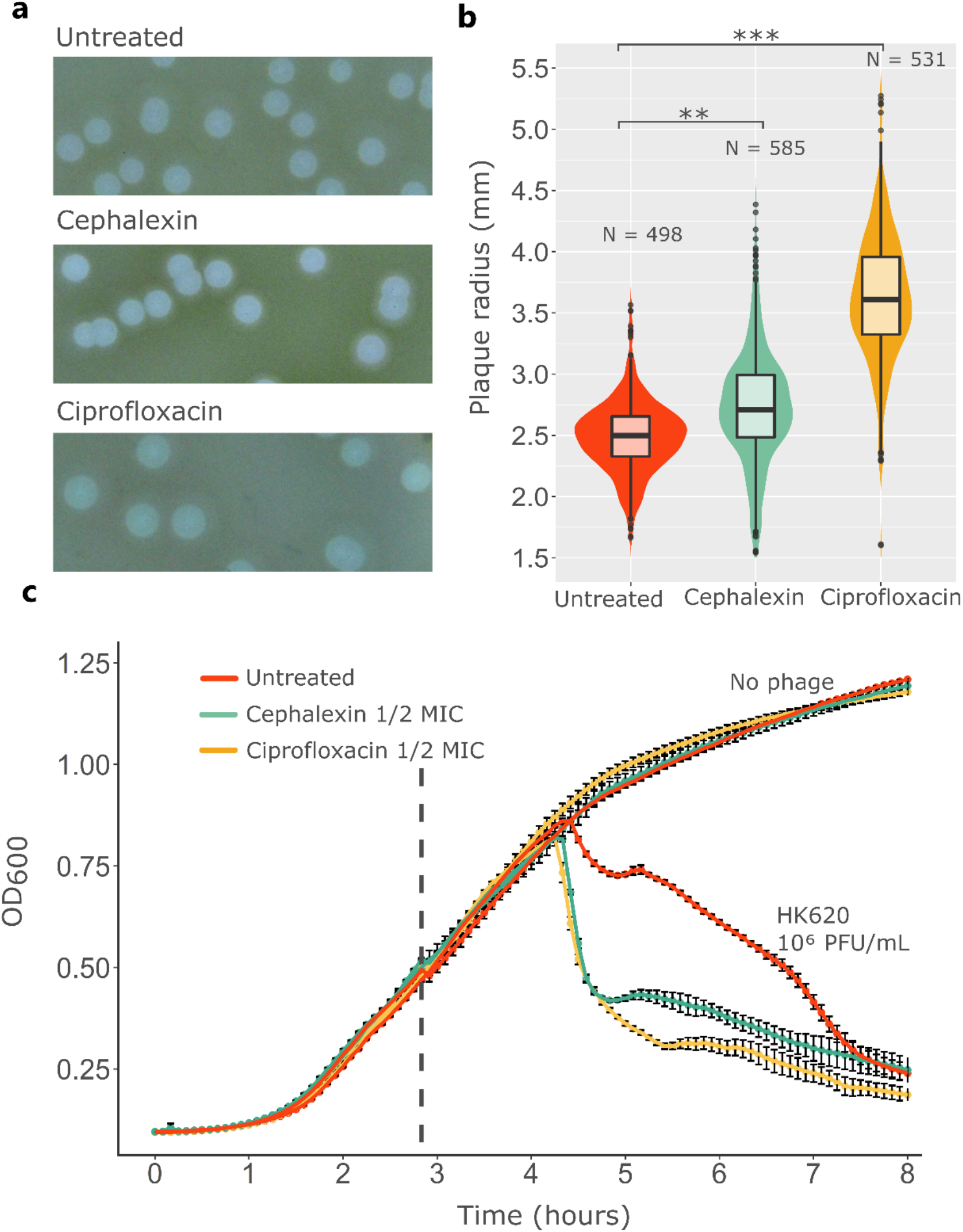
Phage-Antibiotic Synergy (PAS) of phage HK620 and filamentation-inducing antibiotics. **(a)** HK620 lysis plaques from a double agar overlay assay on a lawn of *E. coli* TD2158 PL4, with and without antibiotics at ½ MIC. **(b)** Lysis plaque radii comparison between treatments. P values for a two-tailed test of less than 0.001 are summarised with three asterisks, less than 0.01 are summarised with two asterisks, and non-significant differences are marked as “ns”. **(c)** PAS in liquid culture. The antibiotics were present at inoculation time. The vertical dashed line represents the time of phage HK620 addition at a concentration of 10^6^ PFU/mL. Error bars indicate standard error of mean (s.e.m.).

### Heterogeneous response to ciprofloxacin and cephalexin creates a subpopulation of filamentous bacteria

One of the landmarks of fluoroquinolone treatment is the induction of bacterial filamentation due to the SOS response activation [21]. β-lactam antibiotics can also cause filamentation, although through a different mechanism, which relies on the disruption of peptidoglycan metabolism via the inhibition of bacterial penicillin binding proteins (PBPs) [22]. In both cases, the number of bacteria that filament and the average cell length is dose-dependent. After testing several concentrations of each antibiotic, we decided to keep ½ of the MIC as our experimental concentration, since it led to the largest synergistic effect on phage propagation. In the presence of such concentrations of ciprofloxacin or cephalexin, a significant subpopulation of *E. coli* TD2158 PL4 fails to divide properly and produce filaments of variable sizes (Fig. S1a, b). Whereas 99% of the untreated population displayed a cell length below 5.2 μm, 30% and 58% of the population was above this size in the presence of ciprofloxacin and cephalexin, respectively, with the tail of the cell length distribution extending up to 12 μm. In both cases, the filaments are composed of a single, continuous cytoplasm harbouring several chromosomes (Fig. S1c).

### Single-phage tracking of infection

It was therefore important to track infection at the single cell and single phage scale to account for the heterogeneity of the bacterial population and to detect any tropism of phages for a given subpopulation. To study how phage HK620 interacts with each of the morphologically-different *E. coli* subpopulations, we tracked phage adsorption and infection at single-cell level using epifluorescence microscopy. Phage HK620 was transiently labelled with GFP after propagating it in the TD2158 PL4 host producing the plasmid-encoded protein fusion HkbS-GFP. Labelled phages do not encode the fusion on their own genome, allowing to properly observe only a single round of phage adsorption. As HkbS is the major capsid protein (MCP) of phage HK620, we needed to avoid any steric hindrance that could prevent correct virion assembly and stability, we thus introduced a 15-amino acid linker between the MCP and the GFP sequences (see Methods section). For image analysis, we separated elongated from non-elongated subpopulations, the length threshold used to discriminate between regular-size and filamentous cells was set to 5.2 μm since 99 % of the untreated *E. coli* TD2158 PL4 cells in exponential growth phase fall below this threshold (Fig. S1).

Tracking of phage virions by fluorescence microscopy showed that mean HK620 adsorption per cell increased after ciprofloxacin or cephalexin treatment compared to the untreated condition (Fig. 2b). As previously theorised [23] it can be observed that phage adsorption follows a Poisson distribution since it takes place at a constant rate and since an individual HK620 adsorption event does not modify the likelihood of further phage adsorption. Under antibiotic treatment, 48.4% of the ciprofloxacin and 63.0% of the cephalexin-treated cells had at least one phage attached to them versus 34.2% of the untreated culture (Fig. 2b). The presence of at least one phage adsorbed to the cell surface was considered as an indicator of potential infection, since adsorption constitutes the first step of phage replication. To look closer at the effects of antibiotic-induced filamentation, we separated each antibiotic-treated population in two subpopulations according to their length (Fig. 2c). This allowed us to evaluate the effect of cell length heterogeneity on phage adsorption: 37.7% and 46.5% of the regular-size cells (< 5.2 μm) in the ciprofloxacin and cephalexin-treated cultures display three or more phages per cell, respectively, whereas these proportions raised up to 58.4% and 67.2% for elongated cells (? 5.2 μm). On average, antibiotic-treated cultures display twice (ciprofloxacin) or three times (cephalexin) more phage adsorbed per cell than the untreated cultures (Fig. 2d). This difference in phage adsorption can be explained through a rather simple mechanism: since subinhibitory ciprofloxacin and cephalexin concentrations do not impair growth in mass but still inhibits cell division (Fig. S2), a net reduction in cell numbers is observed. Thus, if the number of phages remains constant, an indirect consequence of these antibiotic treatments is an increase of phage/bacteria ratio due to cells that fail to divide. Furthermore, if we focus on the antibiotic-treated cultures, filamentous bacteria adsorb on average twice more phage per cell than their regular-size counterparts (Fig. 2e). Together these results show that in a heterogeneous population of bacteria, filaments, whatever the agent causing elongation, adsorb more phages than regular-size cells. Since phage adsorption is proportional to the collision frequency between the phage and its host [24], we hypothesised that the increased level of attachment to filamentous cells was due to their enlarged cell surface. To evaluate this, we calculated the mean HK620 adsorption per unit of surface between subpopulations (Fig. 2f). Conclusively, our results suggest there is no significant difference of adsorption per unit of surface between filaments and regular-sized cells. This supports the idea that phage adsorption is proportional to the bacterial size, making it more frequent in longer cells due to an increased probability of phage-host encounter.

**Figure 2:**
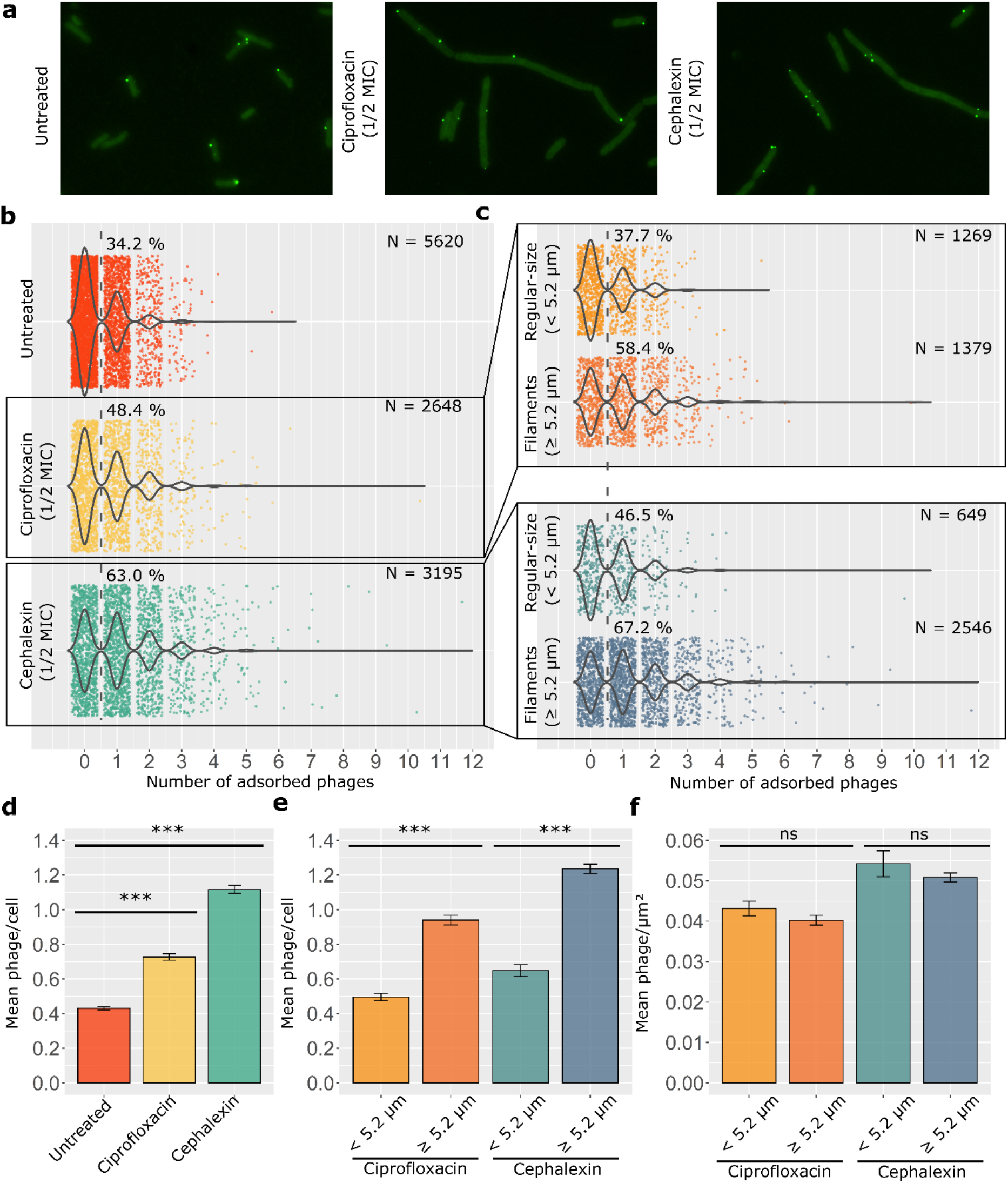
Distribution of HK620 adsorption on *E. coli* TD2158 PL4 cells. **(a)** Epifluorescence microscopy of GFP-coated HK620 on the surface of untreated, ciprofloxacin, and cephalexin-treated *E. coli* cells. **(b)** Comparison of phage adsorption distribution between untreated, ciprofloxacin, and cephalexin-treated cultures; the dashed lanes and percentages indicate the fraction of cells with one or more phages per cell. **(c)** Comparison of frequencies of adsorption between regular-size and filamentous subpopulations within the ciprofloxacin and cephalexin-treated cultures. Bacterial cells were segmented and fluorescent-phage foci were detected and assigned to the corresponding cell using MicrobeJ, an ImageJ plug-in. Percentages represent the fraction of cells that have adsorbed three or more phages on their surface. **(d)** Mean number of phage adsorption for untreated, ciprofloxacin, and cephalexin-treated cultures. **(e)** Mean number of phages adsorbed to regular-size or filamentous subpopulations within the ciprofloxacin and cephalexin-treated cultures. **(f)** Mean number of phage adsorption per unit of surface for regular-size and filamentous subpopulations within the ciprofloxacin and cephalexin-treated cultures. Error bars indicate standard error of mean (s.e.m.). P values less than 0.001 for a two tailed test are summarised with three asterisks, non-significant differences are marked as “ns”.

### Filaments are infected by HK620 significantly more often than regular-size cells

As adsorption is only the first step of a phage productive cycle, we wondered whether the rate of HK620 infection was also different between regular-size and filamenting cells. To evaluate this, we used HK620 *hkcEF::P_rrnB_-gfp* [25], a reporter phage that express GFP fluorescence during phage intracellular replication (Fig. 3a, see Movie 1 and Fig. S4 for the time-lapse analysis of HK620 *hkcEF::P_rrnB_-gfp* infection). The non-compartmentalized GFP fluorescence suggests that filamentous cells share the same cytoplasm and get infected as a single cell. The fraction of infected cells was calculated in the presence and absence of ciprofloxacin or cephalexin and plotted according to MOI (Fig. 3b, c). This MOI was calculated as PFU/CFU at the time of infection. The rate of infected cells observed within the filamentous subpopulation was significantly higher than in regular-sized bacteria. These results are in accordance with the observed heterogeneity in phage adsorption. At low initial MOIs such as 0.5 phage/cell, we measured a larger number of fluorescent filaments than regular-sized cells. This confirms the fact that filaments are more prone to interact with phages, due to their enlarged surfaces, than non-filamentous cells. For example, around 50% of elongated cells were infected at a MOI of 0.5 in the presence of ciprofloxacin, whereas it required a MOI of 1-1.5 to reach the same proportion of infected cells in the regular-sized subpopulation. It is only when phages largely outnumber bacteria (initial MOI ? 5) that most cells got infected no matter their size.

**Figure 3:**
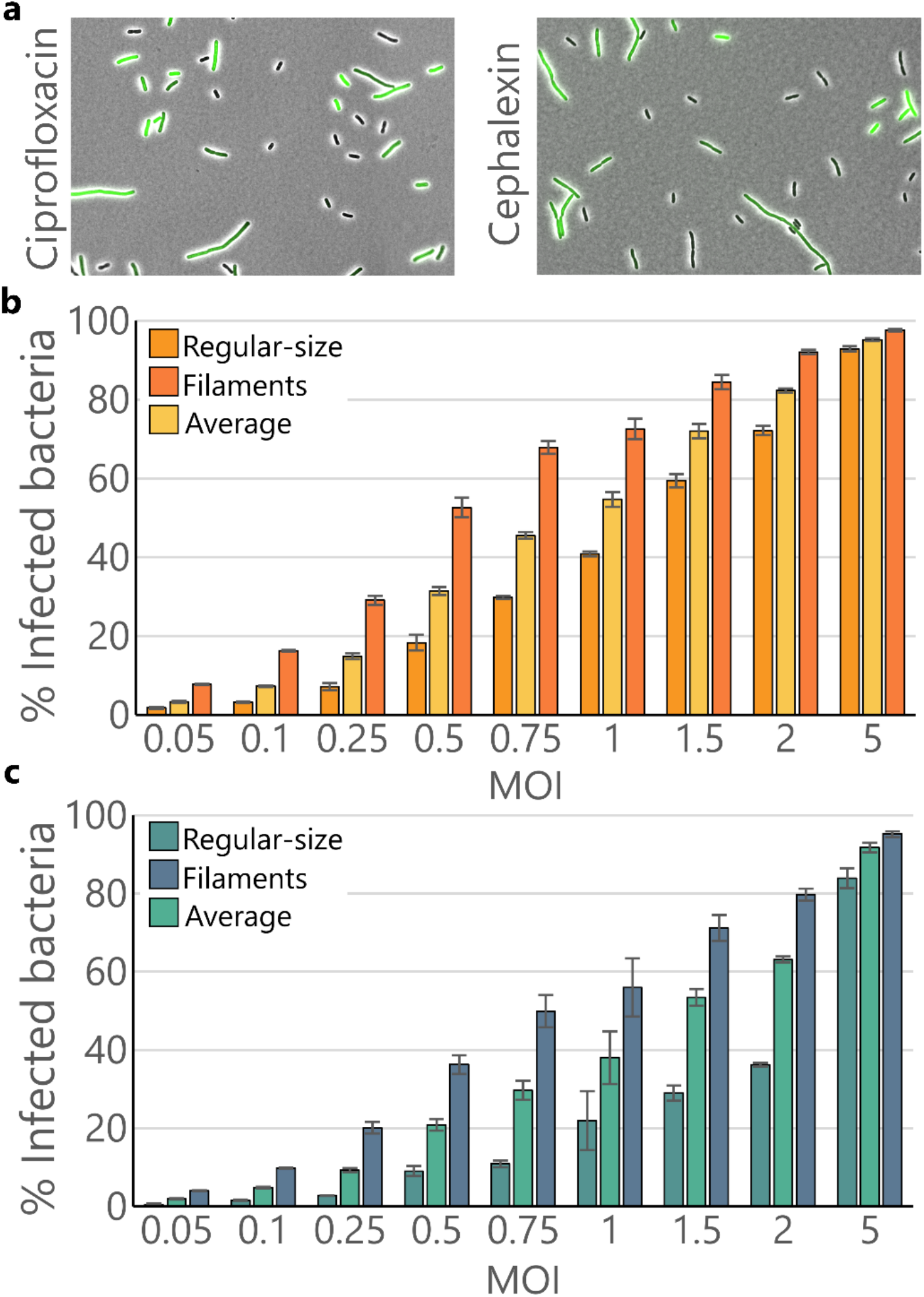
Infection rates of regular-size and filamentous subpopulations within antibiotic-treated cultures. **(a)** Phage-encoded GFP reporter allows to discriminate between naïve and infected cells at time = 20 minutes. Images show HK620 infection on *E. coli* TD2158 PL4 at MOI = 1. Antibiotics are present at ½ of the MIC. **(b)** Infection rate of each subpopulation in ciprofloxacin-treated (½ MIC) cultures. **(c)** Infection rate of each subpopulation in cephalexin-treated (½ MIC) cultures. Error bars indicate standard error of mean (s.e.m.). **Movie 1: Fluorescence increase upon HK620 *hkcEF::PrrnB-gfp* infection.** Time-lapse recording of *E. coli* TD2158 PL4 cells infected with phage HK620 *hkcEF::PrrnB-gfp* at MOI = 1. Recording of the infection started at time = 10 min. after phage addition and proceeded for 45 minutes with capture intervals of 5 minutes.

### Phage HK620 lyse filaments more often than regular-size cells

In order to spot which subpopulation was effectively lysed upon infection, we performed a flow cytometry analysis after a single round of phage infection. An untreated *E. coli* TD2158 culture was analysed and used to create a gating that includes 90% of the population (Fig. 4a). As previously stated, this culture consists mostly of regular-size cells (Fig. S1). Upon sublethal antibiotic treatments, a filamentous subpopulation emerged accounting for 43% and 51% of the total population for cephalexin and ciprofloxacin-treated cultures, respectively (Fig. 4b, c). We then subjected the same antibiotic-treated cultures to a single round of phage lysis before fixation and analysis by flow cytometry. Cytograms showed a neat decrease of the filamentous subpopulations that had appeared upon antibiotic treatments (Fig. 4d, e). These results unambiguously show that longer cells are preferentially lysed during infection.

**Figure 4:**
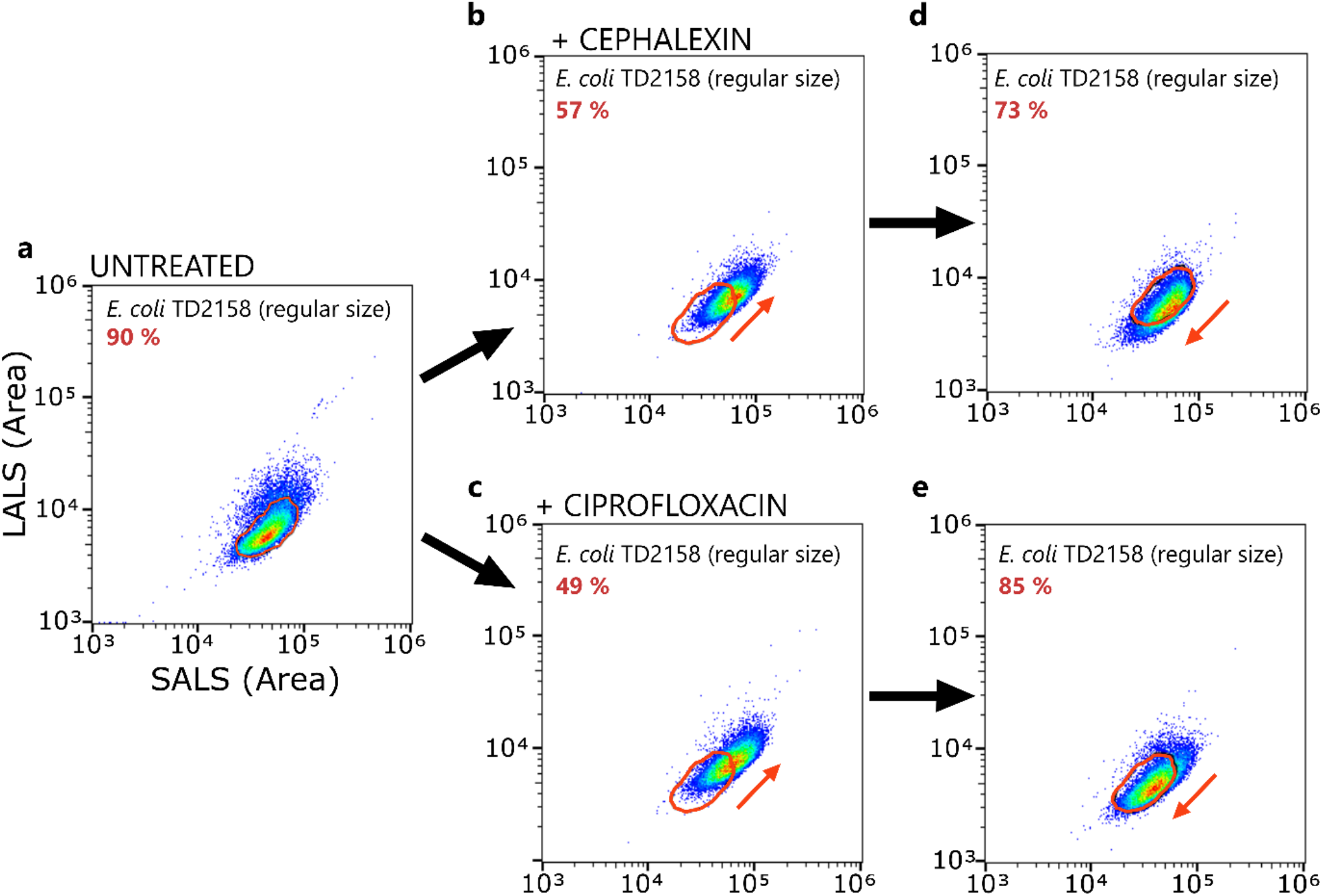
Cytograms of *E. coli* TD2158 PL4 populations with or without phage and antibiotic treatment. **(a)** After 2 h of exponential growth. **(b,c)** After 2 h of exponential growth in the presence of cephalexin and ciprofloxacin at ½ of the MIC. **(d,e)** After one of round of HK620 lysis at MOI = 0.5. Percentages in red represents the population within the gating (red ellipse).

### Cephalexin and ciprofloxacin produce an increase in phage HK620 burst-size

An additional question concerns the influence of the antibiotics on the latent period, which is the time frame between the beginning of infection and the release of the phage’s progeny, and the burst-size, which is the number of phages produced per infected cell. We thus measured HK620 burst-size on untreated cells and compared it to ciprofloxacin or cephalexin-treated conditions using the classical One Step Growth Curve (OSGC) experiment (Fig. S3). This protocol can only be performed in bulk cultures, making impossible to distinguish differences between the previously observed length subpopulations. This experiment showed that phage HK620 burst size increased by 28% and 36% in the presence of ciprofloxacin or cephalexin, respectively. This increase in productivity is likely beneficial for phage propagation, since more phages are produced per lytic cycle. Surprisingly, the larger burst-sizes were not linked to changes in the latent period. These parameters are usually related since a longer latent period might increase the time available for phage intracellular production. However, our results seem to indicate that a larger burst-size might arise from enhanced rate of phage production in the presence of antibiotics, rather than longer assembly periods.

### A mathematical model to understand the synergy between antibiotic and phage treatments

We developed a model to integrate all the effects discussed above on the dynamics of a population of bacteria exposed to a combination of phage and antibiotic treatments (Fig. 5 and SI). Bacteria may be exposed to a sublethal dose of antibiotic at rate σ which blocks the division of bacteria and induces filamentation of susceptible cells (*S* and *F* refer to the densities of regular size and filamentous cells, respectively). Filamentous bacteria may eventually recover from the filamentous state at rate *γ* and resume to normal division. Bacteria may also be exposed to a bacteriophage growing lytically and *V* refers to the density of free viral particles. We show with this model that when we account for the higher adsorption rate on filamentous cells (i.e., when *a_F_* > *a_s_*) we recover the synergistic effect of antibiotic and phage on the total biomass of the bacterial cells (Fig. 5b and SI), which accurately describes the experimental results (Fig. 1, Fig. 2).

**Figure 5.**
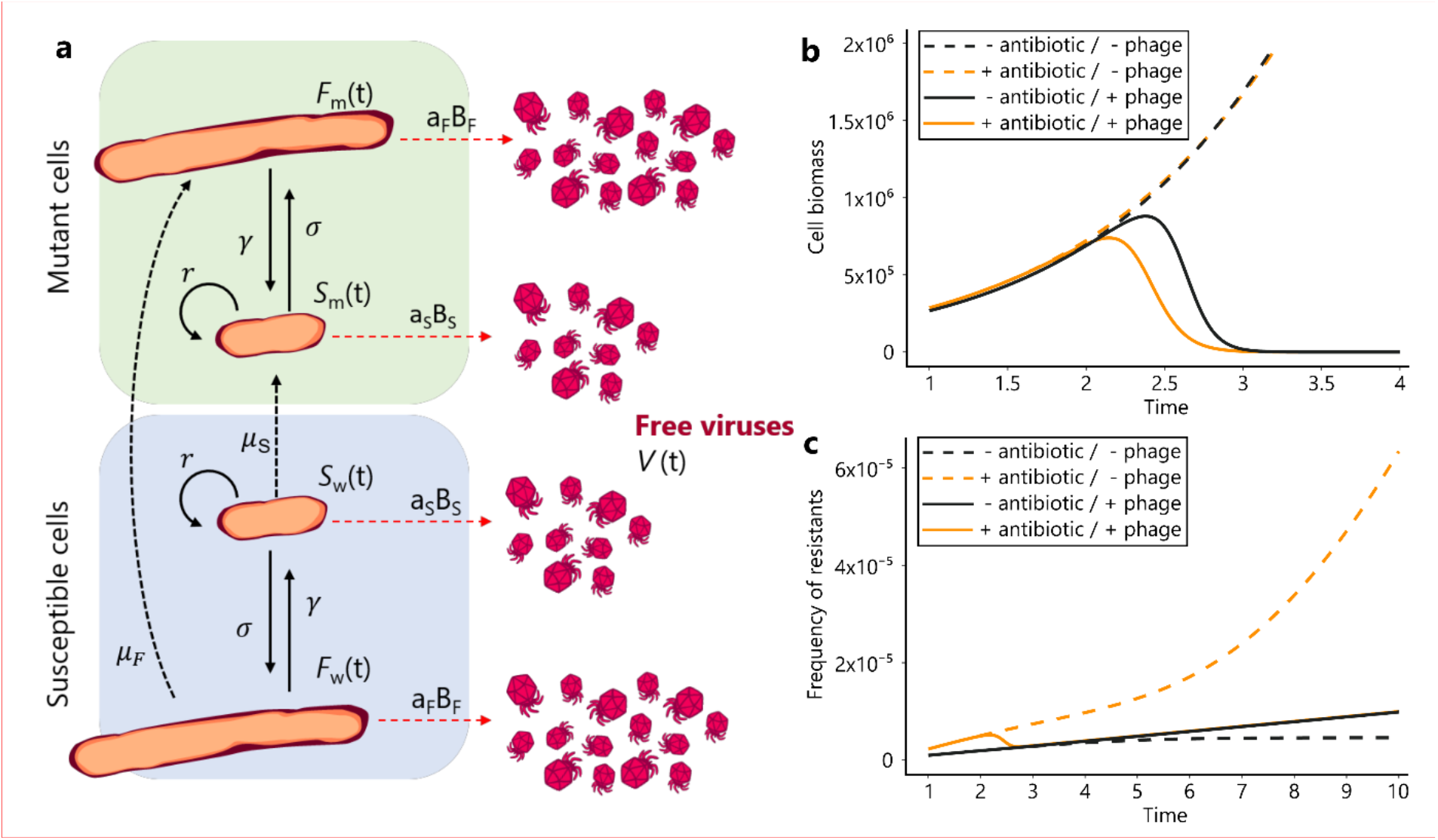
A mathematical model to understand the synergy between antibiotics and phages. **(a)** Schematic representation of the interactions between bacteria and phages in the presence of filamentation-inducing antibiotics (see supplementary information). **(b)** Dynamics of the biomass 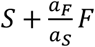 of bacteria in the presence of sublethal doses of antibiotics and/or in the presence of phages. **(c)** Dynamics of the frequency of mutant bacteria in the presence of sublethal doses of antibiotics and/or in the presence of phages. The bottom black dashed line indicates the dynamics of the frequency of mutant bacteria in the absence of antibiotics. Note that we obtain the same dynamics if phages are present because phage only affect this dynamic if some filamenting bacteria are present.

Our model can also be used to explore the effect of phage predation on the accumulation of mutations in populations exposed to sublethal doses of antibiotics. Indeed, filamentous bacteria are known to launch DNA repair systems and thus to exhibit higher mutation rates [26]. Such an increase in mutation rates increases the likelihood that individual cells will acquire mutations that allow them to escape the effects of antibiotic. Yet, if phage predation is preferentially directed towards filamentous cells (i.e., when *a_F_* > *a_s_*) the frequency of filamentous cells drops (see Fig. 4 and SI) which may reduce the accumulation of mutations in the bacteria population. Our model predicts that even if sublethal doses of antibiotic may trigger mutagenesis, the combination of antibiotic and phage should yield lower frequency of resistance mutations (fig. 5c). We test this prediction in the following section.

### Phage infection decreases SOS-induced mutagenesis

As fluoroquinolones are prone to induce mutagenesis through the SOS-response [27], we wondered if phage addition could modify the mutagenesis rate associated with the ciprofloxacin treatment. For this, we adapted the fluctuation test protocol, which estimates the number of rifampicin resistant mutants (Rif^R^) obtained in the presence of a given stress [28] (see the Methods section for details). First, we verified by OD_600_ measurements, that neither antibiotic nor phage treatment significantly impacted bacterial growth at the concentrations and MOIs employed (Fig. S2, S5a). Although HK620 behaves lytically at 37ºC, it remains a temperate phage. To rule out the occurrence of lysogeny under our experimental conditions we looked at the presence of HK620 integrated in the *E. coli* chromosome at the end of the experiment through colony PCR (Fig. S5b). As described recently [29], in the presence of ciprofloxacin lysogeny is highly reduced, since fluoroquinolones are strong inducers of the SOS response, which in many cases can reduce prophage stability. Indeed, 100 CFU obtained from different replicates plated after 8 hours of HK620 infection in the presence of ciprofloxacin were pooled in 5 groups and screened for HK620 integration. No amplification was observed within any of the pooled colonies (Fig. S5c). To evaluate HK620 effect on mutagenesis we measured the number of Rif^R^ colonies that appeared in untreated versus ciprofloxacin-treated cultures after 24 hours. As expected, the mutagenesis rate increased fivefold in the presence of ciprofloxacin, due to error-prone replication occurring upon SOS response activation. We then measured the effect of phage HK620 addition in a ciprofloxacin-treated culture. Strikingly, this rate decreased by over more than five times when HK620 was added in the presence of the antibiotic (Fig. 6). This result suggests that, in agreement with our theoretical prediction (Fig. 5 and SI), while ciprofloxacin increased mutagenesis through the activation of the SOS-response, phage killing seems to reduce this effect. It is remarkable that this effect is achieved even when phages are added in very low doses such as 30 PFU/well (MOI ~10^-7^). To correlate phage infection to SOS-response, we used the *pcda’-gfp* and *pcda’-mcherry* reporter fusions, previously described as an accurate reporter of the SOS-response (Fig. S6) [30]. As shown above, phage predation occurred preferentially within the elongated subpopulation, which constitutes the majority of the SOS-induced subpopulation. Hence, we propose that phages preferentially target a subpopulation under active DNA repair (prone to mutagenesis). Therefore, the combination of phage and antibiotics may carry two benefits: first, a faster bacterial killing, and second a reduction of the influx of antibiotic resistance.

**Figure 6:**
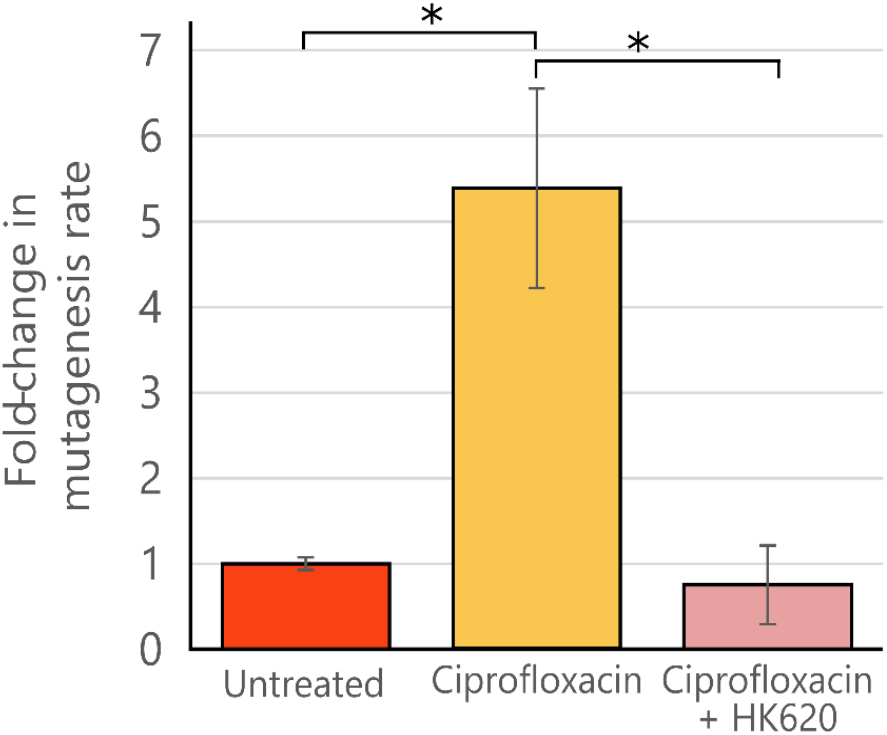
Phage HK620 effect on bacterial mutagenesis. Fold-increase in mutagenesis rate following ciprofloxacin treatment, and its reduction after the addition of HK620 30 PFU/well at 11 hours post-inoculation. P values of less than 0.05 for a one-tailed test are summarised with a single asterisk.

## Discussion

### PAS biological significance

Although phage-antibiotic synergy or PAS has been coined by a relatively recent study [10], it has been observed in multiple labs for several decades. In this work, Comeau *et al.* observed using plaque assays that the bactericidal efficacy increases massively when phages and antibiotics are used in combination. However, PAS has been demonstrated to be highly dependent on the appropriate phage and antibiotic combination, therefore empirical determination has become a rule to determine a particular synergy both *in vitro* and *in vivo* [8,17,18,31]. PAS has been observed on both planktonic and biofilm-embedded bacteria, which thus confers a high potential in human health as well as in animal health and agriculture applications. Several laboratories worldwide are studying PAS, but most studies are restricted to a limited number of combinations, and are rarely focusing on the molecular elucidation of this effect [11–13]. It is also important to elucidate and predict the consequences of such combinatorial treatments from an epidemiological point of view [14]. Importantly, combinatorial treatments not only increase the killing potential of phages and antibiotics but were also shown to be able to re-sensitize antibiotic tolerant bacterial strains in some cases [15–18]. In addition, combination treatments can also induce changes in antibiotic administration habits by decreasing doses, which will reduce selective pressure and thus the potential for the emergence and spread of antibiotic resistance [16].

### Mechanism

Despite the interest of the medical and ecological scientific communities, very few examples in the literature proposed a mechanism for increased cell killing under combined treatments [12,13]. The first proposed mechanism is the delayed lysis hypothesis, which links the burst-size (number of virions produced by a single infected host cell) to the latent period, which is the time between phage adsorption and the release of the first virions [13]. In this work, Kim *et al.* suggested that PAS is the consequence of a prolonged time of particle assembly (latent period) before lysis that increased the yield of virions produced per cell. Host lysis requires the production and accumulation of a phage protein named holin that forms holes in the cell membrane. Some antibiotics, such as β-lactams or those inducing the SOS response, promote cell filamentation but this increase in cell membrane surface does not correlate with an increased phage holin production. This leads to limited amounts of holins to localize in clusters and form holes in the host membrane, hence delaying cell lysis. Strikingly, in a different study it has been shown that under PAS conditions, the lysis time could also be decreased, leading to accelerated lysis giving rise to a higher production of virions [10]. This observation is strengthened by studies using phage T4, where mutations in an anti-holin encoding gene (LIN) lead to a more rapid cell lysis but also produce higher virions yields [32,33]. However, in the present study, although phage HK620 burst-size significantly increased in the presence antibiotics, the lysis time remained unaffected, even though the antibiotics we used are part of the β-lactam family (cephalexin) or induce the SOS response (ciprofloxacin) (Fig. S3). In addition, some studies point to the host membrane fragility in the presence of antibiotics belonging to the β-lactam family that target PBP involved in peptidoglycan synthesis. It was thus proposed that membrane fragility contributed to the decrease of the lysis time [10]. However, this hypothesis would only be valid for β-lactams and does not explain the effect of other classes of antibiotics.

We designed a set of experiments that allowed us to dissect phage infection at the single-phage and single-cell levels and identify the subtle changes at various stages of the phage cycle that could not be identified when looking at bulk experiments. We used filamentation-inducing antibiotics at ½ of the MIC, producing populations of heterogeneous size (Fig. S1). Phage infection was then quantified at the level of phage adsorption step to the bacterial cells (Fig. 2), at the phage genome replication step (Fig. 3) and finally at the lysis step (Fig. 4). All these experiments converge to the same conclusion: elongated cells are infected and lysed by phages more often than regular-sized cells due to their enlarged surfaces. This difference in phage-host interaction was previously overlooked using classical bulk methods such as liquid culture or plaque assay that cannot address bacterial population heterogeneity (Fig. 1, S3). Moreover, compartmentalization of phage-encoded *gfp* expression was not observed in infected filaments (Fig. 3a, Movie 1), suggesting that cells that fail to divide share the same fate after getting infected by a single phage. Such an effect, which can be thought as killing several cells with one virus, can explain the accelerated eradication of bacteria in liquid cultures. In addition, in our experimental settings and using phage HK620 as a model system, the latency period that reflects the time of lysis did not vary according to the presence of antibiotics (Fig. S3). The simplest explanation is that elongated cells catch more phages due to their increased surface (or volume) and that a larger cytoplasm provides more resources to produce and assemble more virions. To date, no technique has been developed yet to determine the burst size of an individual bacterium, and future work should aim at developing such technique using high resolution microscopy. Our results show that under these conditions, the adsorption step is critical and that subsequent infection steps follow on from this. This could not have been determined without the single-cell/phage infection techniques we developed.

### Bacterial resistance emergence

A critical aspect of the combined therapy that needs to be considered is the emergence of bacterial resistance to antibiotics or to phages as a consequence of the selective pressure imposed by the treatments. In the recent years it has been shown that phage genomes rarely encode antibiotic resistance genes, yet the transfer of these genes between bacteria by the mean of phages essentially occurs through generalized transduction, i.e. when a phage particle encapsulates random pieces of the host genome that can carry such resistance genes [34]. On the other hand, antibiotic and phage resistance can occur through mutagenesis that may generate altered versions of a variety of bacterial proteins leading to resistance or modify their production. On the phage-resistance side, such mutations generally affect receptor molecules (lipopolysaccharide or membrane proteins) that become non-permissive to recognition or phage genome ejection [35]. On the antibiotic-resistance side, a variety of mutations can lead to antibiotic resistance according to the antibiotic’s molecular targets (efflux pumps, PBPs, transcription and translation machineries). As an example, mutations in *rpoB,* encoding the β-subunit of the RNA-polymerase, lead to rifampicin resistance, which is the principle used by fluctuation tests to estimate mutagenesis rate [36]. Sublethal concentrations of fluoroquinolones as well as β-lactams tend to increase mutagenesis [27,37]. Our results confirmed that fluoroquinolone-induced elongated cells are SOS-activated and more prone to mutagenesis (Fig. 6, S6). Since this SOS-activated population is a target of choice for phages, we thus wondered if phages could alter the mutagenesis rate. We performed an adapted fluctuation protocol to quantify mutation rates, and our results showed that the addition of phages strongly reduced the mutagenesis induced by ciprofloxacin (Fig. 6). Hence, a combined treatment should reduce *in vivo* resistance acquisition and produce a more efficient therapy than antibiotics or phages applied separately. In addition, it has been shown in a number of recent studies that acquired phage resistance could lead to antibiotics resensitization [35,38–40], reinforcing the need to investigate resistance dynamics to both phage and antibiotics under combined applications.

### Model predictions and experimental validation

As pointed above, the existence of PAS depends on the specific combination of antibiotic and phage. In the present work we focused on antibiotics of two different classes (fluoroquinolone and β-lactam) that both produce cell filamentation (Fig. S1). We contend that the induction of filamentation by sublethal doses of antibiotics may drive PAS. To explore this hypothesis, we developed a mathematical model where the exposition to antibiotic yields filamenting cells. A phage spreading in a bacteria population exposed to sublethal doses of antibiotics will thus have to exploit different types of cell subpopulations. The above experimental results identify the influence of these different cells on the successive steps of the phage life cycle. Our model shows that the higher adsorption rate on filamenting cells affects the composition of the cell population and amplifies the reduction in the population biomass because larger cells are preferentially killed by the phage. Besides, this mechanistic model also captures the influence of phage predation on the reduction in the influx of mutant cells.

Hence, our mechanistic description on the effects of antibiotics on the morphology and the physiology of bacterial cells and their impact on phage dynamics provides a deeper understanding of the effects of combining phage and antibiotic treatments. Combination therapy may provide two distinct beneficial effects. First, a purely demographic effect where combination reduces the biomass of bacterial population (the classical PAS effect). Second, an evolutionary effect where combination reduces the influx of mutations in the bacteria population. Both effects could help maximize the efficacy of phage therapy. Yet, an evaluation of the efficacy of combination therapy needs to account for other processes such as the evolution of resistance to phage and to the antibiotic used in combination. In addition, our model could be readily extended to take into consideration the effects of the immune response of the host on the dynamics of the bacteria [14].

## Materials and methods

### Bacterial strains, phages and culture conditions

Phage HK620 and its host *E. coli* TD2158 were originally provided by J. Clark and S. Barbirz [19,41,42]. Strain TD2158 PL4 was obtained after curation of inducible prophages using mitomycin C [20]. The engineered phage HK620 *hkcEF::P_rrnB_-gfp* was constructed in a previous work [25]. Unless otherwise indicated, bacterial cultures were carried out in Lysogeny Broth (LB) with 1.5% agar for solid media. Phage propagation was performed by infection of a log-phase bacterial culture at 37°C, under 180 rpm agitation until culture clearing. Phages were collected by filtration of the lysate supernatant and further purified by centrifugation [43].

### Minimum inhibitory concentration determination

To determine which antibiotic concentrations impaired significantly *E. coli* TD2158 PL4 growth, OD_600_ curves were performed at 37°C under 180 rpm agitation for 16 hours using a 96-well microplate reader (Spark multimode microplate reader, Tecan). Bacteria were inoculated at OD_600_ = 0.025 in LB medium with ½ serial dilutions of antibiotics. MIC was designed as the lowest concentration with at least 15% of reduction in the area under the growth curve. The maximum antibiotic concentration that allowed undisturbed growth was considered as ½ of the MIC.

### Phage HK620 latent-period determination

In order to set up every further experiment, HK620 latent-period was determined through the One-Step Growth Curve method (Fig. S3). A log-phase *E. coli* TD2158 PL4 culture was infected at MOI < 0.001. After 15 min incubation, 100 μL samples were taken and plated at intervals of 5 minutes until 70 minutes post-infection, and lysis plaques counted.

### Phage-Antibiotic Synergy measurements

PAS measurements in liquid cultures were carried out by OD_600_ measurements in a 96-well microplate. Briefly, a log-phase *E. coli* TD2158 PL4 culture was diluted to a final OD_600_ = 0.025. Subinhibitory antibiotic concentrations were added at this time. Replicates of 200 μL of each condition were dispensed in each well. After two hours of incubation at 37ºC under 180 rpm agitation, the same number of phages (phage titer PF/mL) was added to each replicate and lysis was observed by the reduction in OD_600_. Conversely, PAS measurements in semi-solid cultures were performed using the double agar overlay assay. For this, 10 mL of a log-phase culture of *E. coli* TD2158 PL4 at OD_600_ = 1 were infected with a final titer of 10^3^ PFU/mL of phage HK620 and incubated at 37°C under 180 rpm agitation. Twenty-five minutes post-infection, 100 μL of this culture was mixed with 3 mL of soft-LBA (0.75% agar) and plated over a 20 mL bottom layer of 1.5 % LBA on a petri dish. Antibiotics were added at sublethal concentrations in the bottom layer. Plates containing 100 PFU each were imaged using a digital camera (Nikon D5300). Plaque diameter was measured using a homemade software designed by Leon Espinosa (available upon request) and average plaque diameter was plotted for each condition.

### Antibiotic effect on division time and cell surface

A log-phase *E. coli* TD2158 PL4 culture (OD_600_ = 1) was diluted by 1:1000. Two μL of this dilution were added to an Ibidi μ-Dish 35 mm high microscopy dish (IBIDI, Martinsried, Germany) and squeezed with 1 mm thick 1% agarose-LB pad. Antibiotics were added to the pad according to the previously determined sublethal concentrations. Bacteria were imaged on an inverted phase-contrast microscope (model) thermo-controlled at 37° C. For each condition, three to five individual cells were selected at time = 0 and time-lapse imaged with 5 minutes intervals for 3 h. Accurate bacterial segmentation was carried out using the Misic tool [44]. For each condition, cell number and surface fold-change were plotted over time.

### GFP-coated HK620

To avoid stability problems related to a genomic fusion of GFP to HK620 major capsid protein HkbS, a transient GFP labelling was carried out by propagating the phage on an *E. coli* TD2158 PL4 strain carrying the fusion encoded by plasmid *pUC18-PrplU-hkbS-gfp.* This strain provides a GFP labelled HkbS that incorporates randomly into the capsid during phage assembly together with the unlabelled phage-encoded HkbS. Since the fusion is not phage-encoded, it allows to monitor adsorption without further HkbS-GFP synthesis once the genome is injected. To obtain GFP-labelled phages cells were grown to OD_600_ = 1 and infected with 10^6^ PFU/mL of phage HK620 at 37° C, 180 rpm. After complete lysis (3-4 hours) cell debris were pelleted by centrifugation at 4000 rpm and the supernatant filtered through a 0.2 μm syringe filter to keep only the labelled phage fraction. Homogenous fluorescent labelling of HK620 was obtained using this protocol.

### Single-cell infection

An overnight culture of *E. coli* TD2158 PL4 was diluted to final OD_600_ = 0.025 in 10 mL of LB medium, and supplemented with the appropriate sublethal antibiotic concentrations when required. At OD_600_ ≈ 0.8, phages were added to reach a final MOI between 0.1 to 10. Thirty minutes post-infection aliquots were sampled and fixed by diluting 1:1 in PBS buffer PFA 4% solution to stop phage replication and prevent cell lysis. For adsorption analysis, unabsorbed phages were washed by centrifugation followed by resuspension of the pellet in PBS-PFA 2% solution. For all samples, the mix was put on a coverslip, gently squeezed under a 1 mm thick 1% agarose pad, and directly imaged on an inverted epifluorescence microscope (Nikon TiE) using an oil immersion 100X NA 1.45 objective. Images were acquired using a cooled camera (Hamamatsu Orca Fusion). Acquisition was carried out using Nikon’s NIS-Element software.

### SOS response induction reporter

To compare the level of SOS activation between regular-size cells and filaments, *E. coli* TD2158 PL4 cells harbouring the SOS activation reporter plasmid *pcda’-gfp* (Norman et al., 2005) were grown in the presence of increasing ciprofloxacin concentrations, ranging from 2.5 to 20 ng/mL. Two hours post-inoculation (mid-log phase) cells were fixed in 1:1 PBS buffer 4% PFA and observed under the microscope. Morphological parameters and GFP fluorescence intensity were measured for each individual bacterium. To evaluate the frequency of HK620 *hkcEF::P_rrnB_-gfp* infection in SOS-triggered cells the *gfp* reporter gene in *pcda’-gfp* was replaced by *mcherry* using the SphI/HindII sites flanking the *gfp* gene. Two hours post-inoculation in the presence of ciprofloxacin (½ MIC), *E. coli* TD2158 PL4 /*pcda’-mcherry* cells were infected with HK620 *hkcEF::P_rrnB_-gfp* at MOI = 1. Infection was interrupted after 30 minutes by diluting the samples 1:1 in PBS buffer PFA 4%. The SOS reporter *(pcda’-mcherry)* and HK620 *hkcEF::P_rrnB_-gfp* infection were imaged using the red and green fluorescence channels, respectively.

### Image analysis

Cell image analysis was performed using MicrobeJ [45]. Cell shape parameters were directly measured from phase-contrast microscopy. For phage adsorption quantification, automated foci detection was carried out using the maxima foci function of MicrobeJ, after background subtraction and thresholding the GFP channel to isolate individual fluorescent phages. To quantify phage infection of HK620 *hkcEF::P_rrnB_-gfp* the integrated fluorescence of each cell was measured in the GFP. Statistical analysis was conducted in R [46] and figures were produced using the package ggplot2 [47]. To calculate average phage adsorption per unit of surface, *E. coli* shape was considered as a cylinder with two half spheres on each extremity. Bacterial length (*L*), measured from pole to pole, and average width (*d*) was determined for each bacterium using the image analysis software MicrobeJ. We then used these measurements to approximate bacterial cell surface through the following equation: *π*d*(L-d) + 4*π*(d/2)^2^,* The first term represents the surface of the cylinder of length *(L-d,* and the second, the surface of the two hemispheres of radius (*d*//2) at each pole.

### Susceptibility to phage lysis through flow-cytometry

A log-phase culture of *E. coli* TD2158 PL4 carrying plasmid pP_rpIU_-*gfp* was infected with HK620 at MOIs ranging from 0.1 to 1. Lysis was allowed to take place only once and the infection was stopped 60 min post-infection by fixing the culture with an equal volume of paraformaldehyde 4% solution in PBS 1X. Samples were analysed using the compact flow cytometer A50-micro (Apogee Flow Systems, UK) equipped with an argon ion laser (Asbly, wavelength excitation 488 nm, 50 mV) and specific fluorescence filter set (Green (Gn): 535/35 nm, Orange (Or): 585/20 nm and Red (Rd): >610 nm). Calibration beads (Apogee Flow Systems, 1 μm, excitation 488 nm and a broad fluorescence emission in the Gn, Or and Rd channels, 5,000 event/μl^-1^) were used as a standard. Each sample was run in triplicate. Data were acquired in Log scale using PC control v3.40 and histogram v110.0 softwares (Apogee Flow Systems) and analysed with FlowJoV10 software (TreeStarInc).

### Mutation rates measurements

The frequency of Rif^R^ CFU after 20 hours of growth was determined as follows. A log-phase, OD_600_ = 1 culture was diluted 1:10^6^ (final concentration 200-500 CFU/mL) and 100 μL were dispensed in each well of a U-shaped bottom 96-well microplate (Nunclon Delta-Treated, U-Shaped-Bottom Microplate, Nunc). Antibiotics, if used, were added at ½ of the MIC at this point. The plate was covered with a sealing tape and incubated at 37 °C, 180 rpm, in a humidity cassette to minimise culture evaporation. After 20 h of incubation, the total volume (100 μL) of 84 wells was plated onto separate LBA plates supplemented with 75 μg/mL rifampicin. The remaining 12 wells were serially diluted and plated on non-selective plates to count the total number of CFU. Mutation rates were estimated by the MSS-MLE algorithm provided by the FALCOR calculator (https://lianglab.brocku.ca/FALCOR/). For the experiments in which we assessed phage effect in Rif^R^ mutant frequencies, HK620 phages were added in all wells at 11 h post inoculation with a final titer of 4×10^3^ PFU/well. Each condition was repeated at least three times.

## Acknowledgments

We are thankful to all members of the Phages@LCB group for stimulating discussions and suggestions and particularly to Aurélia Battesti and Nicolas Ginet for critical reading and improvement of the manuscript. We thank Alice Boulanger for designing the HkbS-GFP fusion. This work was supported by two exchange grants from CNRS and the Shota Rustaveli National Science Foundation that allowed I.P. stays in our lab. The Mission for Transversal and Interdisciplinary Initiatives (MITI) from CNRS supported this work through the allowance of a doctoral fellowship to M.A.’s group, as well as a collaborative grant “Adaptation of the living to its environment” to S.G.. The Centre National pour la Recherche Scientifique (CNRS), Aix-Marseille Université (AMU) and the Institut de Microbiologie de la Méditerranée (IMM) all support our research and provide on campus exceptional facilities.

---

**Figure S1: Effect of subinhibitory antibiotic concentrations on *E. coli* TD2158 PL4 morphology. (a)** Phase-contrast microscopy images of exponential growing *E. coli* after 2 hours post-inoculation in the presence of filamentation-inducing antibiotics. **(b)** Cell-length distribution of *E. coli* population under different treatments. Mean cell length was of 2.85 μm, 5.16 μm, and 7.59 μm for the untreated, ciprofloxacin-treated and cephalexin-treated samples, respectively. Percentages represent the filamentous subpopulation, here considered with a length equal or higher than 5.2 μm. P values of less than 0.001 for a two tailed test are summarised with three asterisks. **(c)** DAPI staining of each treatment showing the distribution of the bacterial nucleoids within the cytoplasm.

**Figure S2: Tracking of *E. coli* TD2158 PL4 growth and division rates through phase-contrast microscopy. (a)** Average number of fully-segmented cells in an *E. coli* microcolony over time at 37°C, starting from a single cell. Average generation times were of 17:50 min, 20:25 min and 21:30 min for the untreated, ciprofloxacin and cephalexin conditions respectively. **(b)** Average surface fold-change in the same microcolonies. **(c)** Binary mask of the fully segmented microcolonies obtained through MiSiC.

**Figure S3: One-Step Growth Curves of phage HK620 on *E. coli* TD2158 PL4 at 37 °C.** HK620 displays a latent period of 40 min and a burst-size of approximately 400 PFU per infected TD2158 PL4 cell. Ciprofloxacin and cephalexin did not significantly modify the latent period, but produced an increased burst size of 28% and 36%, respectively.

**Figure S4: GFP-fluorescence build-up in HK620 *hkcEF::P_rrnB_-gfp* infected cells.** Comparison of fluorescence intensity fold-change over time between infected and uninfected cells in movie 1. *E. coli* TD2158 PL4 and phage HK620 *hkcEF::P_rrnB_-gfp* were mixed at MOI = 1 at time = 0 minutes. Intensity was measured for N > 15 bacteria belonging to each group. Vertical-dashed lines represent the lysis of the fluorescent cells.

**Figure S5: HK620 behaves fully lytic in the presence of ciprofloxacin. (a)** TD2158 PL4 growth curves untreated, in the presence of ciprofloxacin (½ MIC), or with both ciprofloxacin (½ MIC) and phage HK620 (30 PFU/well). The dashed line represents the time of phage addition (time = 11 hours). **(b)** Schematic representation of primer design to screen for HK620 integration. If the prophage is present, a fragment of 264 bp will be amplified. **(c)** The resulting colony-PCR on pooled clones recovered after 9 hours of infection (time = 20 hours) revealing the absence of the integrated phage (lanes 1 to 5) and a positive control of a TD2158 PL4 HK620 lysogen (lane 6). M = molecular weight marker.

**Figure S6: SOS activation in regular-size cells and filaments**. **(a)** The *pcda’-gfp* fusion expression was monitored at increasing concentrations of ciprofloxacin in filamentous as well as in non-filamentous subpopulations. The figure shows the percentage of each subpopulation above a threshold defined by the levels of fluorescence found in a ciprofloxacin-free bacterial population. **(b)** Fluorescence foldchange in cells carrying the *pcda’-mcherry* reporter *versus* cell length treated with ciprofloxacin (½ MIC). Percentages in the figure represent the abundance of each subpopulation in the culture. As expected from **(a)**, a strong correlation between cell-length and SOS activation is observed. The dashed vertical line represents the size threshold for filamentation (5.2 μm). Percentages of HK620 *hkcEF::P_rrnB_-gfp* infection on each subpopulation (at MOI=1) were of 41% and 82% for regular-sized cells and filaments, respectively. The filamentous, hypermutagenic subpopulation was infected more frequently than cells retaining a regular shape that had, contrarily, lower SOS activation rates.

