## Supplemental figures and material for "Phage-antibiotic synergy: cell filamentation is a key driver of successful phage predation"

**Supporting Information**

**Figures S1 to S6**

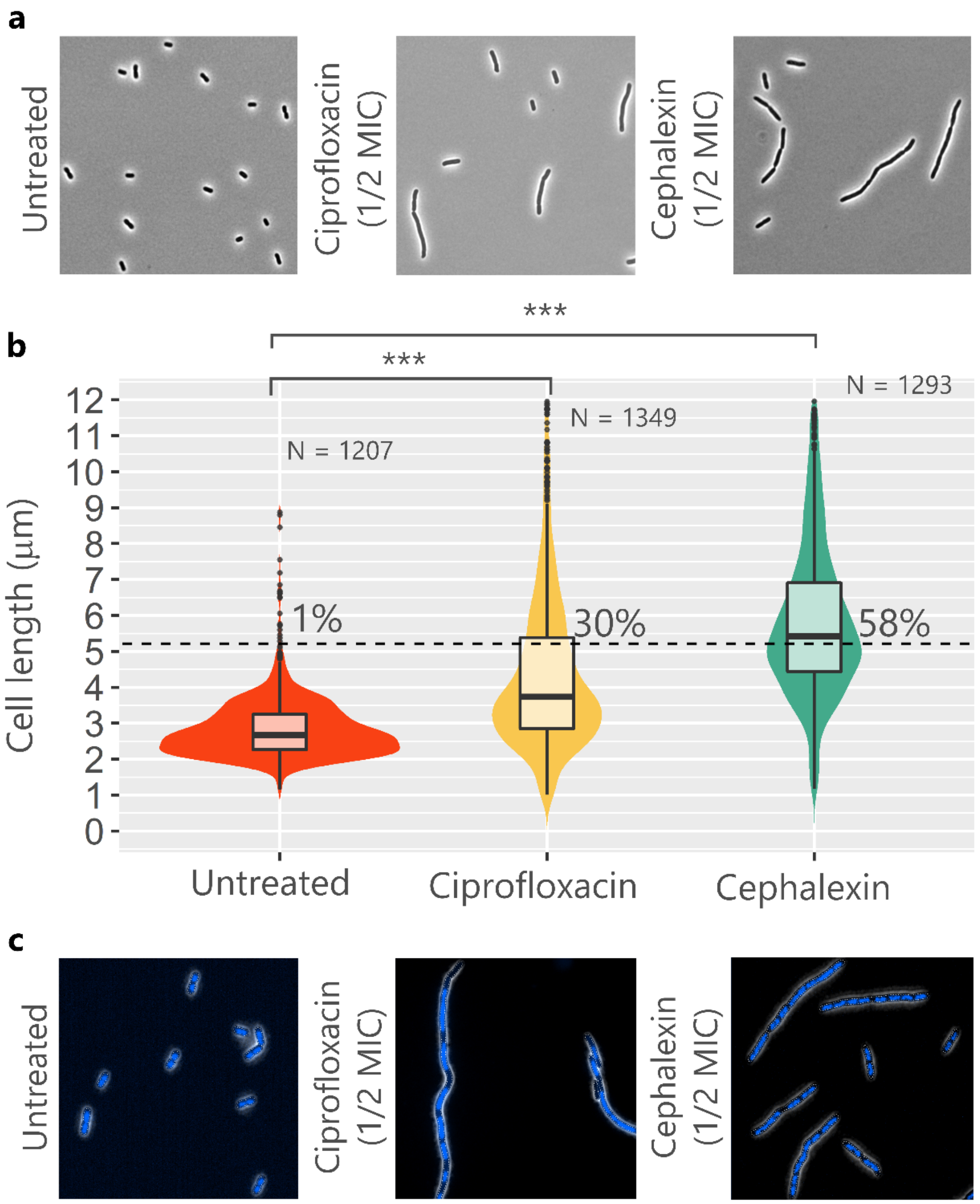


**Figure S1: Effect of subinhibitory antibiotic concentrations on *E. coli* TD2158 PL4 morphology. (a)** Phase-contrast microscopy images of exponential growing *E. coli* after 2 hours post-inoculation in the presence of filamentation-inducing antibiotics. **(b)** Cell-length distribution of *E. coli* population under different treatments. Mean cell length was of 2.85 µm, 5.16 µm, and 7.59 µm for the untreated, ciprofloxacin-treated and cephalexin-treated samples, respectively. Percentages represent the filamentous subpopulation, here considered with a length equal or higher than 5.2 µm. P values of less than 0.001 for a two tailed test are summarised with three asterisks. **(c)** DAPI staining of each treatment showing the distribution of the bacterial nucleoids within the cytoplasm.

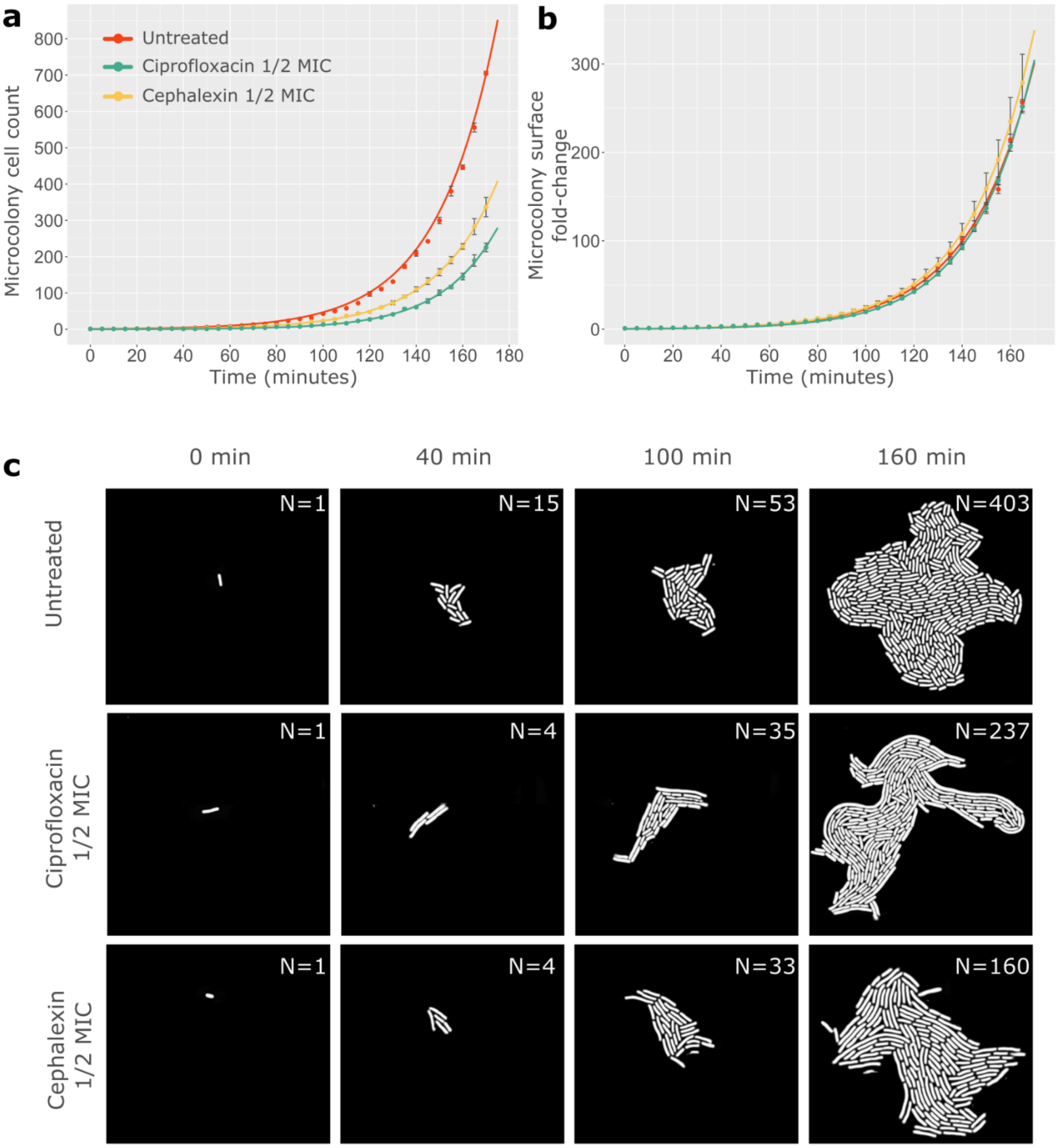


**Figure S2: Tracking of *E. coli* TD2158 PL4 growth and division rates through phase-contrast microscopy. (a)** Average number of fully-segmented cells in an *E. coli* microcolony over time at 37°C, starting from a single cell. Average generation times were of 17:50 min, 20:25 min and 21:30 min for the untreated, ciprofloxacin and cephalexin conditions respectively. **(b)** Average surface fold-change in the same microcolonies. **(c)** Binary mask of the fully segmented microcolonies obtained through MiSiC.

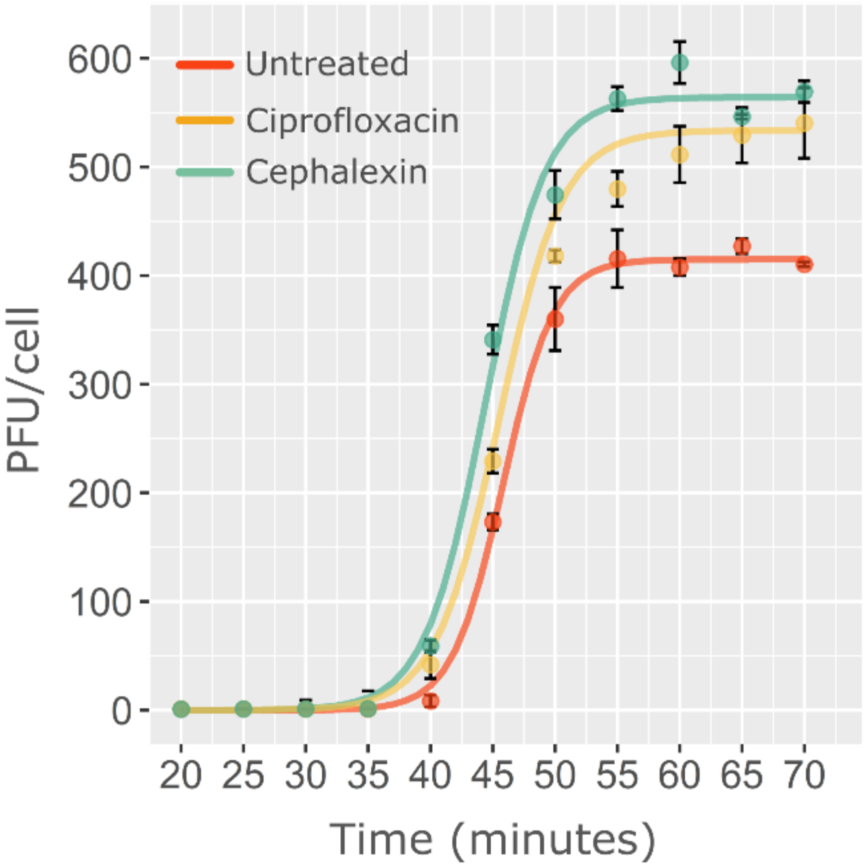


**Figure S3: One-Step Growth Curves of phage HK620 on *E. coli* TD2158 PL4 at 37 °C.** HK620 displays a latent period of 40 min and a burst-size of approximately 400 PFU per infected TD2158 PL4 cell. Ciprofloxacin and cephalexin did not significantly modify the latent period, but produced an increased burst size of 28% and 36%, respectively.

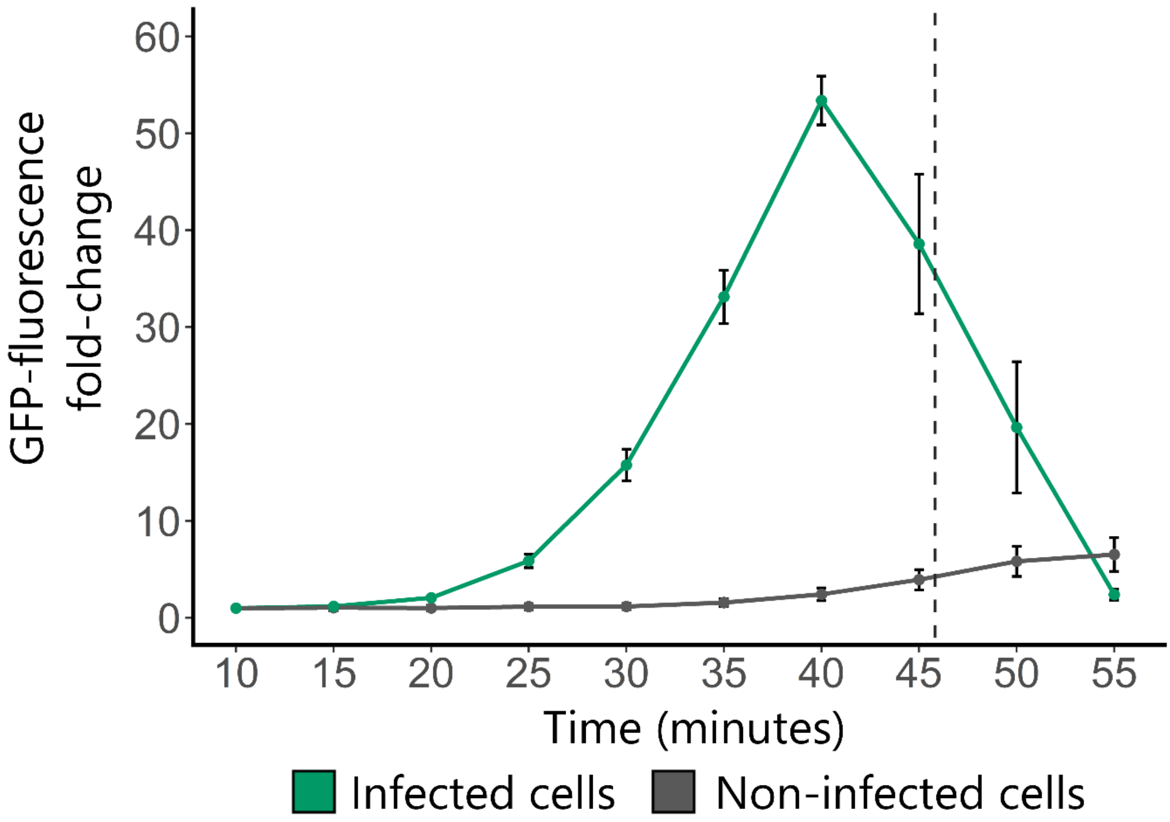


**Figure S4: GFP-fluorescence build-up in HK620 *hkcEF::P$_{rrn}$B-gfp* infected cells.** Comparison of fluorescence intensity fold-change over time between infected and uninfected cells in movie 1. *E. coli* TD2158 PL4 and phage HK620 *hkcEF::P$_{rrn}$B-gfp* were mixed at MOI = 1 at time = 0 minutes. Intensity was measured for N > 15 bacteria belonging to each group. Vertical-dashed lines represent the lysis of the fluorescent cells.

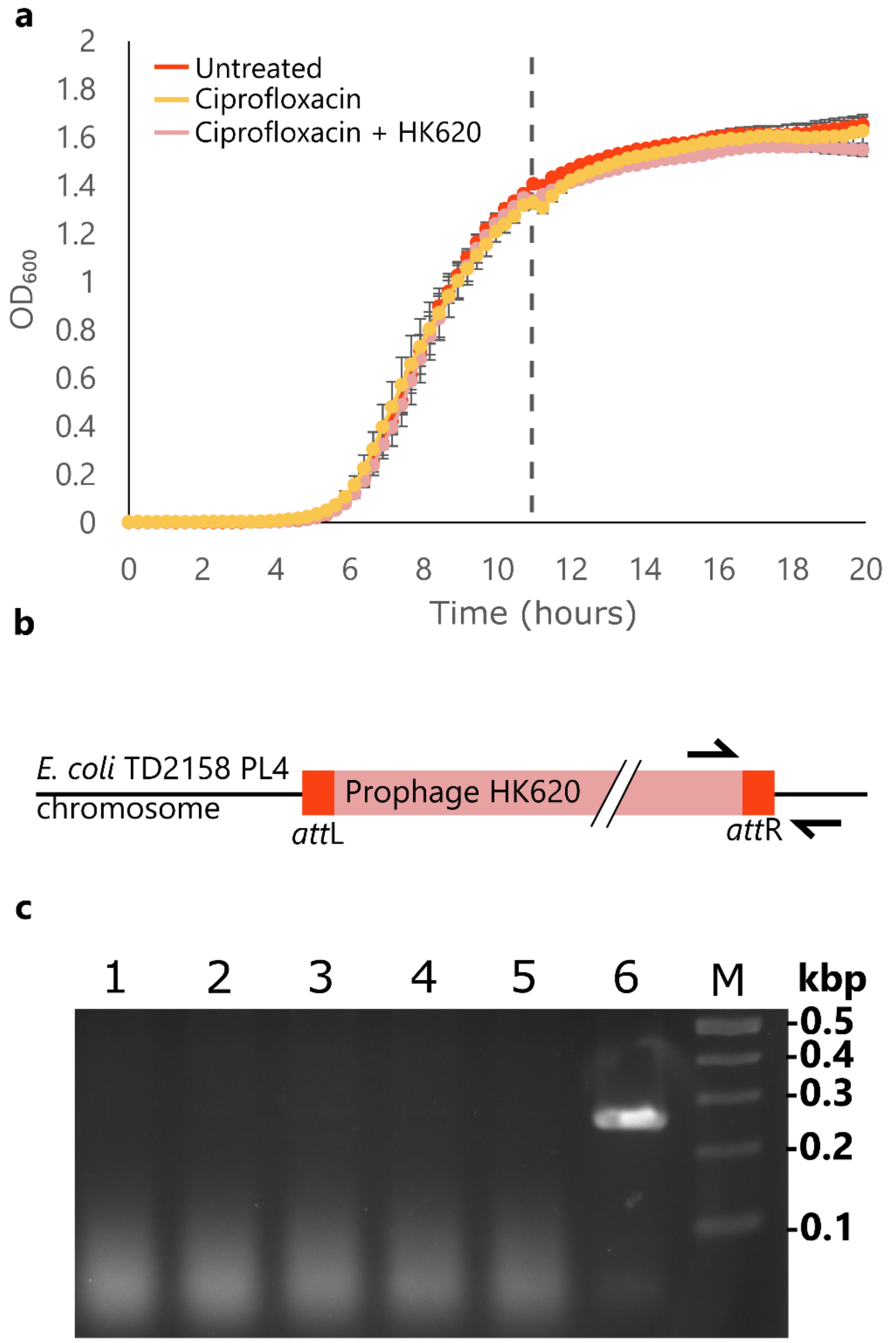


**Figure S5: HK620 behaves fully lytic in the presence of ciprofloxacin. (a)** TD2158 PL4 growth curves untreated, in the presence of ciprofloxacin (½ MIC), or with both ciprofloxacin (½ MIC) and phage HK620 (30 PFU/well). The dashed line represents the time of phage addition (time = 11 hours). **(b)** Schematic representation of primer design to screen for HK620 integration. If the prophage is present, a fragment of 264 bp will be amplified. **(c)** The resulting colony-PCR on pooled clones recovered after 9 hours of infection (time = 20 hours) revealing the absence of the integrated phage (lanes 1 to 5) and a positive control of a TD2158 PL4 HK620 lysogen (lane 6). M = molecular weight marker.

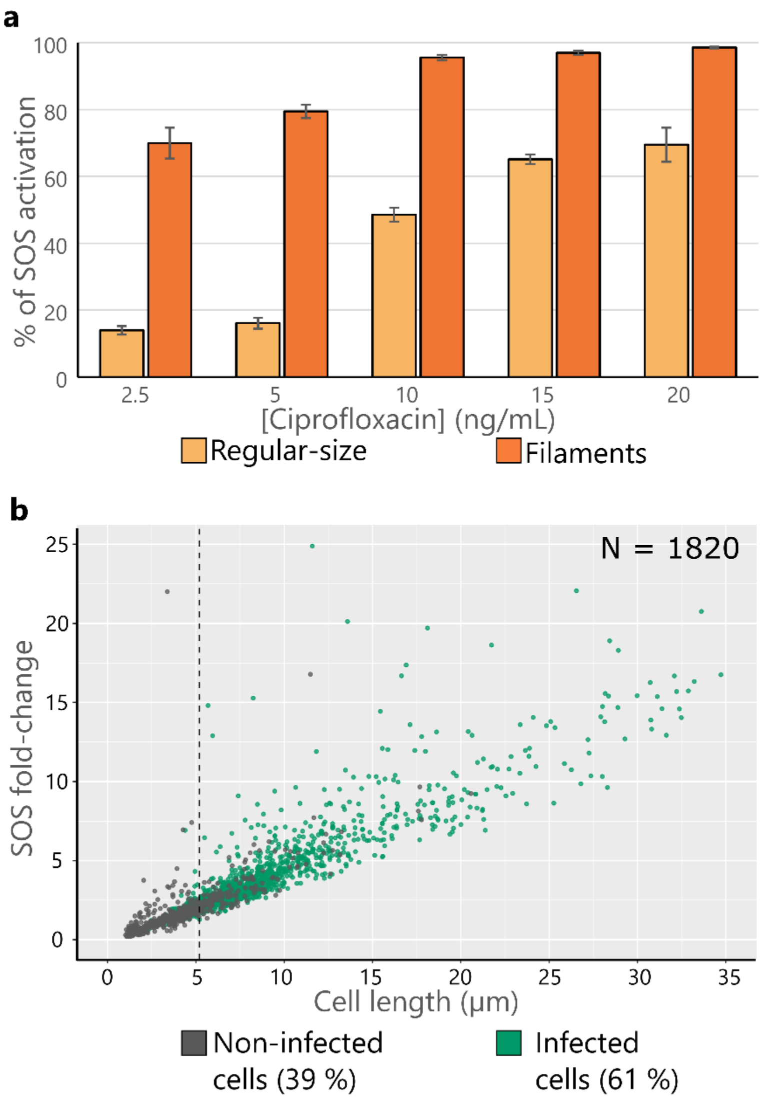


**Figure S6: SOS activation in regular-size cells and filaments**. **(a)** The p*cda'-gfp* fusion expression was monitored at increasing concentrations of ciprofloxacin in filamentous as well as in non-filamentous subpopulations. The figure shows the percentage of each subpopulation above a threshold defined by the levels of fluorescence found in a ciprofloxacin-free bacterial population. **(b)** Fluorescence fold-change in cells carrying the p*cda'-mcherry* reporter *versus* cell length treated with ciprofloxacin (½ MIC). Percentages in the figure represent the abundance of each subpopulation in the culture. As expected from **(a)**, a strong correlation between cell-length and SOS activation is observed. The dashed vertical line represents the size threshold for filamentation (5.2 µm). Percentages of HK620 *hkcEF::*$P_{rrnB}$-*gfp* infection on each subpopulation (at MOI=1) were of 41% and 82% for regular-sized cells and filaments, respectively. The filamentous, hypermutagenic subpopulation was infected more frequently than cells retaining a regular shape that had, contrarily, lower SOS activation rates.

### Supporting text

We model the transient dynamics of a population of bacteria subject to a combination of phage and antibiotic treatments. The density of the wild-type bacteria is noted $S_w$. In the absence of antibiotic these cells have a "normal size" (i.e. no filamentation) and reproduce at a density-dependant rate $r\left(1-\frac{N}{K}\right)$ where $r$ is the maximal growth rate, $N$ is total population size of the bacteria and $K$ is the carrying capacity of the bacteria population (i.e. the maximal cell density). Bacteria are assumed to die at a constant rate $d$.

Bacteria may be exposed to a sublethal dose of antibiotic ($\sigma$ measures the rate of exposition to antibiotics) which does not kill the bacteria but induces filamentation ($F_w$ is the density of filamenting bacteria). Filamenting bacteria stop reproducing but we assume they can recover from the filamentous state and recover a regular size at rate $\gamma$. Filamenting bacteria are known to activate DNA repair systems and thus to exhibit higher mutation rates (Gutierrez et al. 2013, Bos et al. 2015). We track the accumulation of mutant bacteria that acquired resistance to other antibiotics. The variables that refer to the densities of these mutant bacteria are indicated with a subscript $m$ while we use the subscript $w$ for wild-type cells. These mutant bacteria emerge from "normal size" cells at rate $\mu_S$ and from filamentous cells at a higher rate $\mu_F$ to account for their higher mutation rate (*i.e.* $\mu_F > \mu_S$). Mutant bacteria are assumed to reproduce and die at the same rate as the wild-type cells.

Bacteria may also be exposed to a virulent phage which kills the infected cells ($V$ is the density of free viral particles). The phage life cycle starts with the adsorption of the viral particle to the $S$ and $F$ cells at rates $a_S$ and $a_F$, respectively. The lysis of $S$ and $F$ infected cells release $B_S$ and $B_F$ of new viral particles, respectively. These viral particles adsorb to new bacteria or die at a constant rate $d_v$. This yields the following dynamical system (see Fig. 5a):

$$\begin{aligned}
\frac{dS_w}{dt} &= r(1-\mu_S)S_w\left(1-\frac{N}{K}\right)+\gamma F_w-(d+a_S\,V+\sigma)S_w \\
\frac{dS_m}{dt} &= r(\mu_S S_w+S_m)\left(1-\frac{N}{K}\right)+\gamma F_m-(d+a_S\,V+\sigma)S_m \\
\frac{dF_w}{dt} &= \sigma S_w-(d+a_F\,V+\gamma+\mu_F)F_w \\
\frac{dF_m}{dt} &= \sigma S_m+\mu_F F_w-(d+a_F\,V+\gamma)F_m \\
\frac{dV}{dt} &= \big(a_S(S_w+S_m)(B_S-1)+a_F(F_w+F_m)(B_F-1)\big)V-d_v V
\end{aligned} \qquad (1)$$

The total bacteria population size is defined as: $N=S+F$ with $S=S_w+S_m$ and $F=F_w+F_m$.

### 1. Demography

First, we focus on the transitory effects of antibiotics and phages on the density of bacteria. In other words, we ignore the mutations and focus on the dynamics of $S$ and $F$ which yields:

$$\frac{dS}{dt} = rS\left(1-\frac{N}{K}\right) + \gamma F - (d + a_S\,V + \sigma)S$$
$$\frac{dF}{dt} = \sigma S - (d + a_F\,V + \gamma)F \qquad (2)$$
$$\frac{dN}{dt} = rS\left(1-\frac{N}{K}\right) - d\,N - \,V(a_S S + a_F F)$$

The final term of the last equation captures the effect of viruses on the bacterial population (see Fig. 1c). We can use our model to simulate the change in the biomass $S + \frac{a_F}{a_S}F$ of cells which accounts for the increased size of the filamenting bacteria which is also responsible for the increases adsorption rate of the phage. This yields Figure 5b which captures the synergy between antibiotics and phages (compare Fig. 5b and Fig. 1c).

This effect depends on the densities of $S$ and $F$ cells. The dynamics of the frequency $f_F = \frac{F}{N}$ of filamentous cells is given by:

$$\frac{df_F}{dt} = \underbrace{\sigma(1-f_F)}_{antibiotic} - \underbrace{\gamma f_F}_{recovery} \underbrace{- r f_F(1-f_F)\left(1-\frac{N}{K}\right)}_{no\ reproduction} - \underbrace{V f_F(1-f_F)(a_F - a_S)}_{bacteriophage} \qquad (3)$$

This equation captures the effect of antibiotics on the proportion of filamentous cells (i.e. higher concentration of antibiotics increases the proportion of filamentous cells). This equation also shows that a higher density of viruses can reduce the proportion of filamentous cells when $a_F > a_S$. The reduction of the proportion of filamentous cells when cultures are exposed to bacteriophages (see Fig. 4) indicates that $a_F > a_S$.

### 2. Evolution

Second, we focus on another indirect benefit of combination therapy emerging from the reduction of the influx of mutations in the bacteria population. To better understand the transient evolutionary dynamics of resistance we focus on dynamics of the frequency of antibiotic resistance $f_m = \frac{S_m+F_m}{N}$ which yields:

$$\frac{df_m}{dt} = r\left(1-\frac{N}{K}\right)\underbrace{\mu_S f_w^S f_S}_{\substack{mutation \\ from\, S}} + \underbrace{\mu_F f_w^F f_F}_{\substack{mutation \\ from\, F}} + \underbrace{r\left(1-\frac{N}{K}\right)(f_m^S - f_m)f_S}_{growth} + \underbrace{V f_F (a_F - a_S)(f_m - f_m^F)}_{bacteriophage} \quad (4)$$

where: $f_m^S = \frac{S_m}{S}$, $f_m^F = \frac{F_m}{F}$, $f_w^S = \frac{S_w}{S}$, $f_w^F = \frac{F_w}{F}$.

The first two terms in equation (4) account for the effects of mutation rates. In particular, the second term captures the increased mutation rate in filamentous cells. The final two terms depend on the distribution of the mutant between the $S$ and the $F$ cells.

To analyse the build-up of this distribution of mutants we can track the frequency of mutations in the two compartments which yields:

$$\begin{aligned} \frac{df_m^S}{dt} &= r\left(\mu_S(1-f_m^S)\right)\left(1-\frac{N}{K}\right) + (f_m^F - f_m^S)\gamma\frac{F}{S} \\ \frac{df_m^F}{dt} &= \mu_F(1-f_m^F) + \sigma\frac{S}{F}(f_m^S - f_m^F) \end{aligned} \quad (5)$$

Let us define $\Delta = (f_m^F - f_m^S)$:

$$\frac{d\Delta}{dt} = \mu_F(1-f_m^F) - r\left(\mu_S(1-f_m^S)\right)\left(1-\frac{N}{K}\right) - \Delta\left(\sigma\frac{S}{F} + \gamma\frac{F}{S}\right)$$

The dynamics of the above equation is driven by the first term when $\mu_F \gg \mu_S$ and implies that $\Delta > 0$ and thus that $f_m^F > f_m^S$. Indeed, the frequency of the mutant increases in the $F$ cells before diffusing in the $S$ cells. This means that both the third and the fourth terms in (4) are negative and tend to slow down the increase in $f_m$ induced by the high mutation rate $\mu_F$ in $F$ cells.

Increasing phage density acts on the change in mean frequency via the fourth term in (4) (higher phage density decreases the fourth term) but it also acts via the frequency of $F$ cells (see equation (3)). Decreasing the density of $F$ cells reduces the influx of new mutations and reduces dramatically the frequency of mutations. We illustrate the effect of phages on antibiotic-induced bacterial mutagenesis in Figure 5c.

### 3. Parameters

| Parameter | Value | Description |
|---|---|---|
| $K$ | $10^7$ $cells$ | Carrying capacity of the bacteria population |
| $\boldsymbol{\mu_S}$ | $10^{-7}$ $cells^{-1}h^{-1}$ | Mutation rate of regular-size cells |
| $\boldsymbol{\mu_F}$ | $5\ 10^{-6}h^{-1}$ | Mutagenesis rate of filaments |
| $r$ | $1\ h^{-1}$ | Bacterial reproduction rate |

| | | |
|---|---|---|
| $d$ | $0\ h^{-1}$ | Mortality rate of cells |
| $d_v$ | $0\ h^{-1}$ | Mortality rate of the free virus |
| $a_S$ | $10^{-7}\ h^{-1}\ cells^{-1}$ | Phage adsorption rate (regular-size cells) |
| $a_F$ | $2.8\ 10^{-7} h^{-1}\ cells^{-1}$ | Phage adsorption rate (filamenting cells) |
| $B_S$ | $100$ | Phage burst-size (regular-size cells) |
| $B_F$ | $200$ | Phage burst-size (filamenting cells) |
| $\gamma$ | $0.01\ h^{-1}$ | Recovery rate from filamentation |
| $\sigma$ | $0.1\ h^{-1}$ | Filamentation rate (induced by antibiotics) |
| $S_w(0)$ | $10^5\ cells$ | Initial density of bacteria (it was assumed that all other cells were initially absent) |
| $V(0)$ | $1000\ virions$ | Initial density of the virus |

**Table S1.** List of parameters used in the mathematical model of PAS and their definition. The values indicated in the tables were used to obtain the curves in Fig. 5b and Fig. 5c.
